# Higher-order co-mutation interactions in mitochondrial genomes

**DOI:** 10.1101/2023.02.13.528359

**Authors:** Rahul K Verma, Pramod Shinde, Ankit Mishra, Sarika Jalan

## Abstract

Pair-wise co-mutation networks of the mitochondrial genome have already provided ample evidences about the roles of genetic interactions in the manifestation of phenotype under altered environmental conditions. Here, we present a method to construct and analyze higher-order interactions, namely, 3-uniform hypergraphs of the mitochondrial genome for different altitude populations to decipher the role of co-mutating variable sites beyond pair-wise interactions. While the weights distribution of such gene hyperedges manifested power-law for all the altitudes, we identified altitude-specific genes based on gene hyperedge weight. This framework of hypergraphs serves a promising avenue for future investigation of nuclear genomes in context of phenotypic association and genetic disorders.

## Introduction

Networks equips us with a statistical approach to model real-world complex systems based on underlying interaction networks [1]. For instance, in cellular activities, biomolecules interact to perform diverse functions which can be studied through protein-protein interactions [2], gene-transcription factor interactions [3], and metabolic interactions [4] networks. Using the network framework, biologists have analyzed complex sub-cellular interactions to access various structural and evolutionary functionality of cells [5]. Such an interaction analysis under the lens of network theory has assisted in deducing biomarkers and designing new experiments [6, 7]. With an advent in network theory concepts, many other details of real-world systems, such as edge weights emphasising on relative importance of interactions, directed edges [8] indicating flow of signal through edges [9], multi-layer structure [10, 11], and evolution of network properties with time aka temporal networks [12, 13] got incorporated while constructing the corresponding model networks.

All these studies relied in describing complex systems through pairs of interacting units (pair-wise interactions), and demonstrated that properties and evolution of real-world complex systems indeed can be modeled as a net of interacting units. Nevertheless, quest for more accurate representations of properties of real-world interactions has continuously been revealing many exciting facets, particularly, as in real-world systems, the interactions occur in much more detailed fashion. Many factors affect the very nature of interactions themselves. One of the most exciting revelations of these pursuits is discovery of higher-order interactions, which emphasize on existence and importance of interactions beyond pair-wise in real-world complex systems. There are shred of evidences which indicate significance of higher-order interactions in governing properties of many real-world complex systems such as ecological [14], biological [15], neuronal [16], and social [17]. In ecological systems, more than two species interact together and affect the ecological dynamics of the ecosystem. In collaboration networks, more than two authors appear in the same article [18]. In epistatic networks, more than two variable sites interact/co-occur for phenotypic benefits of the species, and reactions involve more than two biochemicals to perform a cellular function. The prediction of critical genes or nodes based on graphs assume that two adjacent nodes have similar functional contribution or where we miss the information of other nodes contributing in that module (functional module). The representation of networks as hypergraphs is a spontaneous way of overcoming this assumption.

Thus, a *d*−uniform hypergraph defines simultaneous interactions between *d* nodes forming edges *E*^(*d*)^ = {*v*_1_, *v*_2_, *v*_3_, …, *v*_*d*_}. Hence, 2−uniform hypergraph corresponds to *E*^(2)^ = {*v*_1_, *v*_2_} depicting pair-wise interactions, 3−uniform hypergraph represents triangles with *E*^(3)^ = {*v*_1_, *v*_1_, *v*_3_} and so on. Note that in a hypergraph all the hyperedges need not to have the same size. However, a hypergraph is referred to as a *k*-uniform hypergraph if all hyperedges connect the exactly *k* nodes. Here, we consider a 3-uniform hypergraph where each hyperedge connects three nodes.

Earlier studies on hypergraphs for the brain were based on the temporal co-variation of brain function during learning via cross-link structures [19]. In a similar approach, correlation hypergraphs were formulated based on edges rather than brain regions to analyse topological and functional changes in the brain during neurodevelopment [20]. Higher-order interactions of protein complexes demonstrated the scale-free degree distribution of both the nodes and the hyperedges [21, 22]. Additionally, signalling pathways [23] and protein interactions [24, 25] have been studied under the umbrella of higher-order interactions.

Furthermore, interactions between genetic variations (epistasis) and the genetic background alter the phenotype landscape of the respective population, thereby leading to various studies of mutations among genes [26, 27]. All these investigations are comprised of only pair-wise interactions and hence limit our understanding of genetic evolution [15]. The presence of a third mutation has been displayed to alter the fitness and also modify the way pair-wise interactions occur [28, 29]. An earlier study on higher-order interactions identified a few phenotypes that were not explained by pair-wise interactions while searching for missing heritability in the crossing of two yeast strains for 46 different phenotypes [30]. Another study involving 500 genotypes of yeast tRNA and over 45000 interactions revealed abundance of higher-order interactions which were responsible for dynamics across different genotypes [31]. In another study, higher-order interactions of five different genes were found to affect the complex phenotype in *Saccharomyces cerevisiae* [32]. This and a few other studies investigated the presence and absence of mutations/variations and their effect on the fitness or morphology of the organism. However, Ref. [33] quantified the digenic and transgenic interactions with respect to the fitness of the population by introducing a transgenic score which combined double and triple mutant fitness extracted from the experimental results. Further, the word co-occurrence hypergraphs have been studied to explore the conceptual landscape of mathematics [34].

All these investigations provided sufficient evidence of the existence of higher-order interactions in complex systems and, moreover, highlighted the importance of such interactions and corresponding hypergraphs in steering the evolution and functionality of underlying complex systems. Here, we studied the co-mutation hypergraphs in the human mitochondrial genomes for three different altitude populations, low-altitude (0-500m), middle-altitude (2001-2500m), and high-altitude (*>*4000m). As the altitude increases, the percent of oxygen in the air decreases. This decrease in the oxygen level is more or less 1% for each rise in 500m altitude [35]. We took three populations, two from extremes and one from the middle, to have a comprehensive analysis of mtDNA variations in the framework of hypergraphs. It has been reported earlier that high-altitude leads to a change in mtDNA composition, which might help individuals to adapt better in low-oxygen environments [36]. In this work, we constructed 3-uniform hypergraph based on co-mutation frequency defined for three variable sites simultaneously for all the possible combinations of the available variable sites. Thereafter, we used a two-step threshold selection method to construct the unweighted hypergraph of variable sites and a weighted hypergraph of genes.

In the following texts, the terms triangles and hyperedges were used interchangeably as we considered 3-uniform hypergraphs only, where each hyperedge contained exactly the three nodes. By going one step forward from traditional pair-wise interactions to triangle interactions, we specifically calculated the co-mutation frequency of three variable sites. A two-step threshold selection was used to filter significant hyperedges, one by using a statistical score (*P*_*ijk*_) and the second by using the size of the largest connected component (*N*_*LCC*_). Thereafter, to identify the significant genes, we calculated the weight of each triangle (hyperedge), followed by the mapping of genes yielding gene hypergraph. Then we analyzed the network properties of those genes which participated in higher-order interactions and survived after the application of thresholds for each altitude group. Note that this study considered only 3-uniform hypergraphs. One could consider higher-order uniform hypergraphs or mixed hypergraphs, which would, accordingly, require more sample sizes for statistically reliable results.

## Methods and Material

mtDNA sequences were collected from Ref. [37] for three altitude groups, lowest (0-500m), middle (2001-2500m) and highest (4001m). The ambiguous nucleotides were replaced by the default letter ‘N’ for the calculations. All the sequences were aligned together using Clustal Omega and then mapped to rCRS for gene annotation.

### Construction of higher-order Gene-Gene Interaction (GGI) networks

Here, we construed two types of 3-uniform hypergraphs; the co-mutation hypergraphs with variable sites as nodes and the weighted GGI hypergraphs with genes as nodes (Fig. 2) for each set of the population.

#### Step 1 (Co-mutation Network)

Any position having more than one allele in the samples was considered a variable site. The variable sites were extracted from the aligned sequences for each region separately. For genomic equality, ambiguous nucleotides such as X, M, Y, etc., were replaced with ‘N’ for all the sequences, and tri-allelic sites were not considered.

#### Step 2

To construct a network for each altitude group, nodes were represented by the position of variable sites, and the edges were represented by co-mutation frequency among three nodes (*C*_*ijk*_) defined as

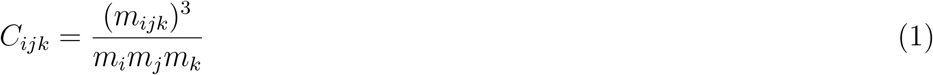

where (*m*_*ijk*_) denotes number of times the minor alleles occur together at *i*^*th*^, *j*^*th*^ and *k*^*th*^ positions, *m*_*i*_, *m*_*j*_ and *m*_*k*_ indicate total number of times the minor allele occurs at *i*^*th*^, *j*^*th*^ and *k*^*th*^ positions, respectively.

#### Step 3 (p−value calculation)

To check the significance of any co-mutation, the threshold has been calculated as

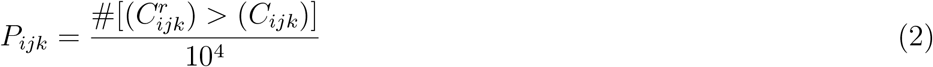

where 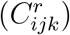 indicates the co-mutation frequency calculated after permuting the alleles at the *i*^*th*^, *j*^*th*^ and *k*^*th*^ positions randomly. 10,000 random simulations were generated and *P*_*ijk*_ was set to 0.05 (standard *p*-value) to consider a co-mutation triangle significance.

#### Step 5 (Higher-order and pair-wise interactions selection)

To define a triangle as filled or unfilled [38], we mapped the variable sites in each triangle (*i, j* and *k*) to their respective genes and if all the genes are different, we considered the corresponding gene triangle as a filled triangle (true triangle) or hyperedge of order 3, otherwise we considered as an unfilled triangle.

#### Step 6 (Co-mutation threshold selection)

To select a co-mutation threshold for the filled triangles, we analyzed the size of the largest connected component (LCC) as a function of *C*_*ijk*_. A threshold was selected based on the change in the size of LCC (Fig. 1).

**Figure 1.**
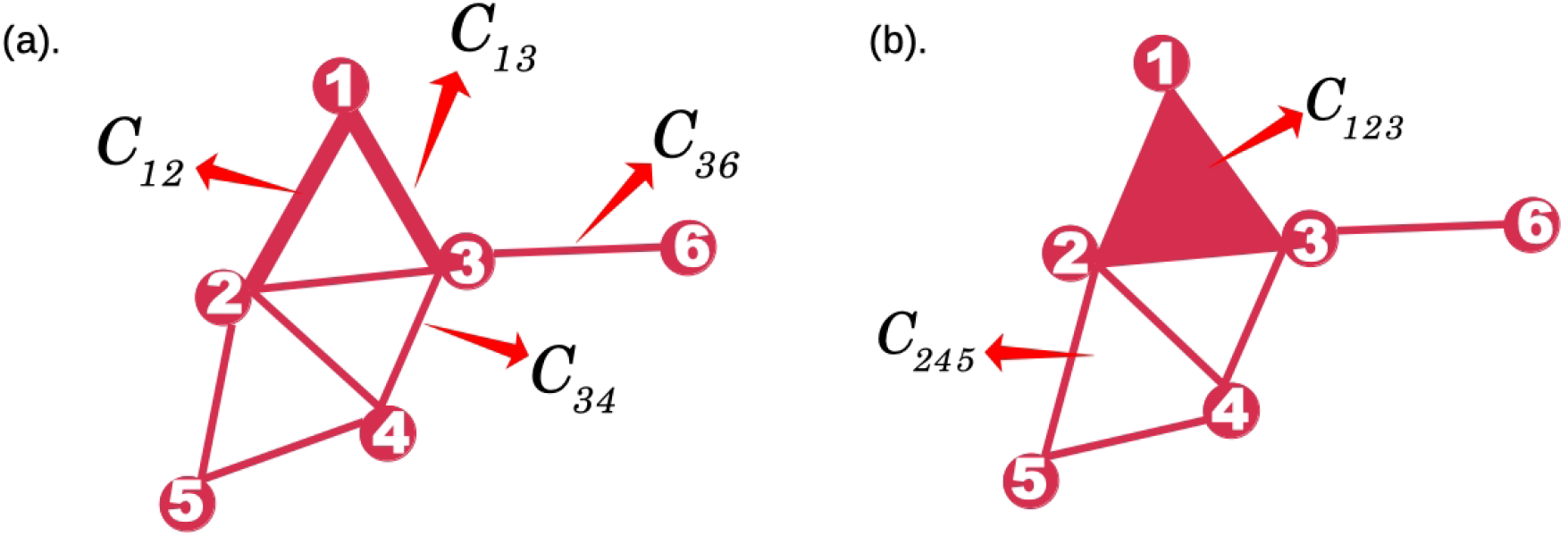
Schematic depiction of pair-wise (left), and triadic (right) interactions based on co-mutation frequency. A thick line (left) and filled triangle (right) indicate that those interactions survived the *p* − *value* test.

#### Hyperdegree of nodes

A hypergraph mathematically represented by *H* = {*V, e*^*H*^} which consists of set of *nodes* and *hyperedges*. The set of *nodes* are represented by *V* = {*v*_1_, *v*_2_, *v*_3_,…, *v*_*N*_} and *hyperedges* by *e*^*H*^ = {*e*_1_, *e*_2_, *e*_3_,…, *e*_*M*_} where *N* and *M* are size of *V* and *e*^*H*^, respectively. Note that, each hyperedge *e*_*α*_, {*α* taking value = 1, 2,…, *M*, **corresponds to** 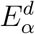 which is a subset of nodes i.e. 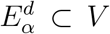. Since, we considered 3−uniform hypergraph, 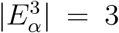 for all *α*. In a hypergraph, the hyperdegree of a node *i* represents the number of hyperedges formed by the node *i*. Next, two types of hypergraphs are constructed in this paper as described in the method section, namely, hyperegraph of the variable sites (*H*_*v*_), and hypergraph of genes (*H*_*g*_). We defined the hyperdegree accordingly, let *d*(*v*)_*n*,2_ and *d*(*v*)_*n*,3_ be the degree of the *n*^*th*^ variable site in the 2-uniform and 3-uniform hypergraphs, respectively. Similarly, let *d*(*g*)_*n*,2_ and *d*(*g*)_*n*,3_ be the degrees of the *n*^*th*^ gene in 2-uniform and 3-uniform hypergraphs, respectively.

### Relative weight of genes

Relative weight (*R*_*n*_) of a gene *n* was calculated based on the number of triangles that gene is participating (which is equal to hyperdegree *d*(*g*)_*n*,3_ of *n*^*th*^ gene) as

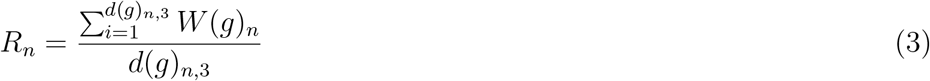

where summation runs for all the triangles in which a given gene *n* is participating, *W*(*g*)_*i*_ is the weight of *i*^*th*^ gene triangle in which gene *n* is participating.

## Results

The following sections analyze the gene hyperedges in terms of their genetic functions, statistical parameters, and associated genetic information. First, we discuss the characteristics of true triangles based on their constituent nodes and hyperedge degrees. Then, we try to incorporate the information from network parameters with the specific haplogroups. Further, we analyse genes involved in each gene triangle to identify the specific genes for each altitude group.

### Characteristics of hyperedges

Initially, after applying the first step filtration based on *P*_*ijk*_, the resultant co-mutation hypergraphs were quite dense, therefore a co-mutation threshold (*C*_*th*_) was further applied to construct a rather sparse network. To identify a value of suitable *C*_*th*_, we analyzed the *N*_*LCC*_ as a function of *C*_*ijk*_ (Fig. 2). An increase in the threshold value would filter out many connections whose weights were smaller than the threshold value. For a threshold value of zero, all the hyperedges would persist. Whereas, for a threshold value of 1, only those hyperedge would persist which have weight exactly equal to 1. We witnessed that there existed a sharp jump in the size of the largest connected cluster as we increased the value of *C*_*th*_. A value just before this value of *C*_*th*_ at which the network becomes segmented was taken as the threshold. This threshold selection method has been defined as network efficiency score [39]. We started with *∼*450 nodes in each altitude group with a potential of 0.1 million combinations and ended up with a very small number of total triangles and true triangles (Table 1). For a given hyperedge, the *C*_*ijk*_ was subjected to the minimal co-mutation frequency value for all three pairs of the triangle. Hence, *C*_*ijk*_ is not a linear combination of pair-wise co-mutation frequencies for all the pairs of *i, j, k* variable sites. After getting the sparse network, we defined the true triangles, such as no variable sites in any hyperedge belong to the same gene for all the hyperedges as at the genetic level, such as a pair of variable sites where both the nodes belong to the same gene would give rise to a self-loop and also transform the triangle into a weighted edge. Hence such connections were also removed. In low-altitude, mid-altitude, and high-altitude *∼*10% of all the triangles were found to be true triangles (Table 1).

**Figure 2.**
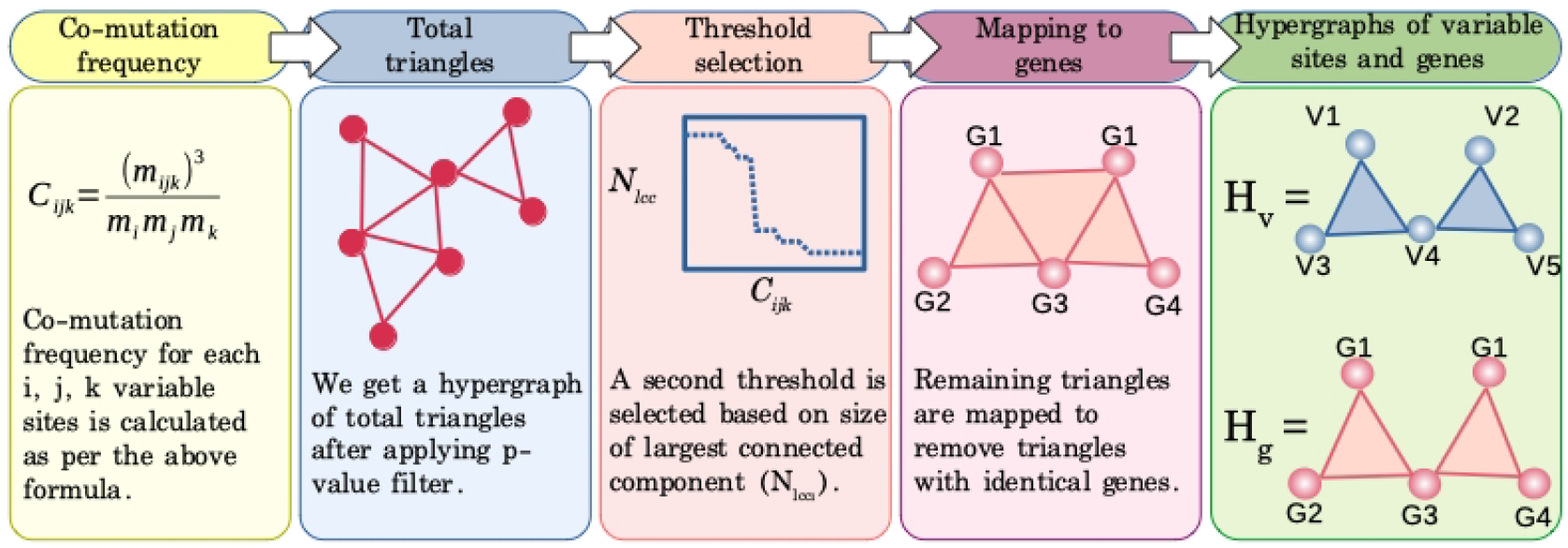
The schematic representation of deducing higher-order interactions from the mutations information of variables sites (a), and construction of co-mutation triangles (b). Co-mutation triangles were then mapped to extract gene triangles (c). The filled gene triangles were identified such that no gene is identical in triangle (d). Finally, a threshold value for higher-order interactions was identified (e) to obtain 3-uniform gene hypergraphs (f).

**Table 1.**
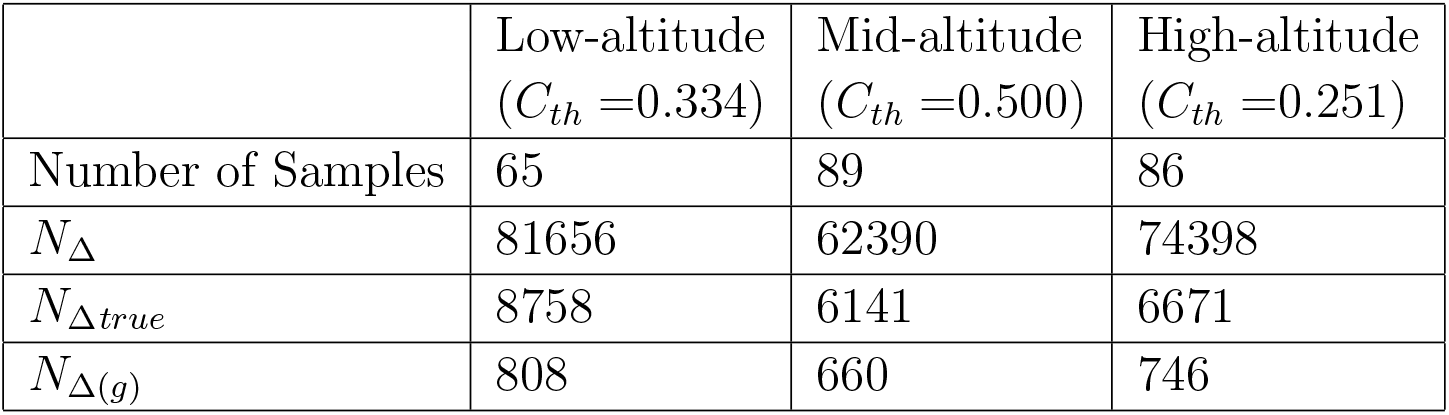
Number of triangles (total number of hyperedges of order 3) before (*N*_Δ_) and after (*N*_Δ*true*_) applying the threshold. The number of gene triangles (*N*_Δ(*g*)_) forming hyperedges (*e*(*g*)_*n*,3_) are considered without counting those in the Control region for the three altitude populations; low-altitude (0 - 500m), mid-altitude (2001 - 2500m) and high-altitude (4001m).

| | Low-altitude<br>($C_{th}=0.334$) | Mid-altitude<br>($C_{th}=0.500$) | High-altitude<br>($C_{th}=0.251$) |
| --- | --- | --- | --- |
| Number of Samples | 65 | 89 | 86 |
| $N_{\Delta}$ | 81656 | 62390 | 74398 |
| $N_{\Delta true}$ | 8758 | 6141 | 6671 |
| $N_{\Delta(g)}$ | 808 | 660 | 746 |

### Variable sites and haplogroups

From this small number of triangles, we checked for the variable sites (nodes) contributing to the formation of triangles in terms of their degree (*d*(*v*)_*n*,3_) (Table 2). In the low-altitude group, node 1598 represented M-haplogroup (M17, M28, M29, M30, M42, M52, B5 and N1). In the mid-altitude group C-haplogroup represented by nodes 14318 (C4) and 15204 (C1, C4, C5 and C7), and in the high-altitude group, K-Haplogroup is represented by nodes 9055 (Z3, K1, K2 and U8), 9698 (K1, K2 and U8), 10550 (K1 and K2) and 11299 (L0, K1, K2 and Y2). It is noted that in the low altitude, only one high-degree node was represented as a haplogroup marker. In contrast, most high-degree nodes in mid and high altitudes represented a few haplogroups. It is to note that despite having very low variable sites in low-altitude, tRNA genes, Trp, Tyr, Gly, and Ser are contributing to the highest node degree. When the difference (Δ*d*(*v*)) of pair-wise (*d*(*v*)_*n*,2_) and triangle degrees (*d*(*v*)_*n*,3_) was plotted, we found that certain variable sites were having large differences compared to most of the nodes. This suggests that a few variable sites show a higher tendency to form hyperedges (Fig. 3 upper panel).

**Table 2.**
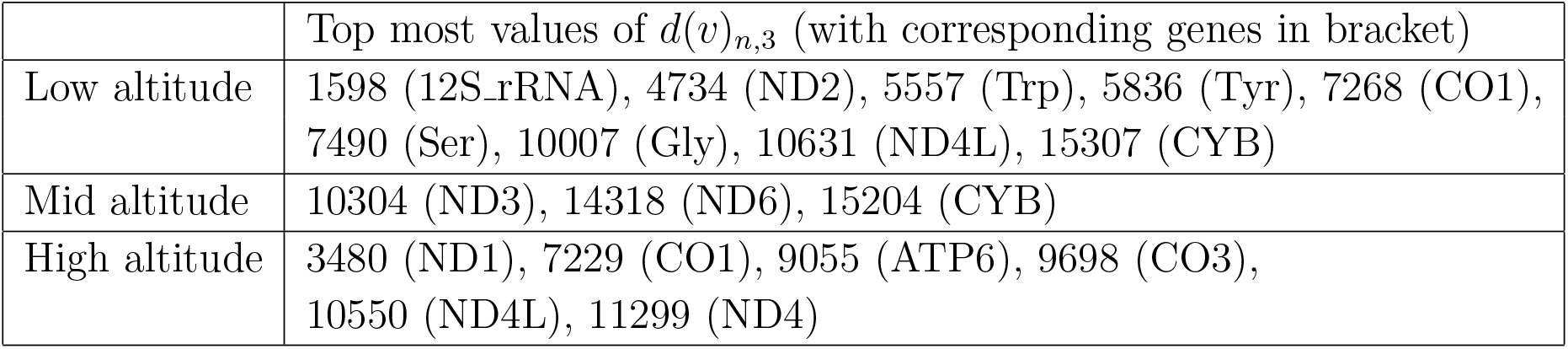
The nodes with the highest contribution in each altitude group.

**Table 3.** Gene triangles with distinct weights in each altitude groups.

|  | Gene triangles with weight |
| --- | --- |
| Low Altitude<br>(0-500m) | ND1-ND2-ND5: 67<br>ATP6-ND2-ND5: 62<br>ATP6-ND1-ND5: 60 |
| Mid Altitude<br>(2001-2500m) | CO1-ND4-ND5: 51<br>CO1-ND1-ND5: 45<br>ND2-ND4-ND5: 43 |
| High altitude<br>(>4001m) | CO1-ND1-ND5: 40<br>ATP6-CYB-ND2: 37<br>CYB-ND1-ND5: 37<br>ATP6-CO1-CYB: 36<br>ATP6-CYB-ND5: 36 |

**Figure 3.**
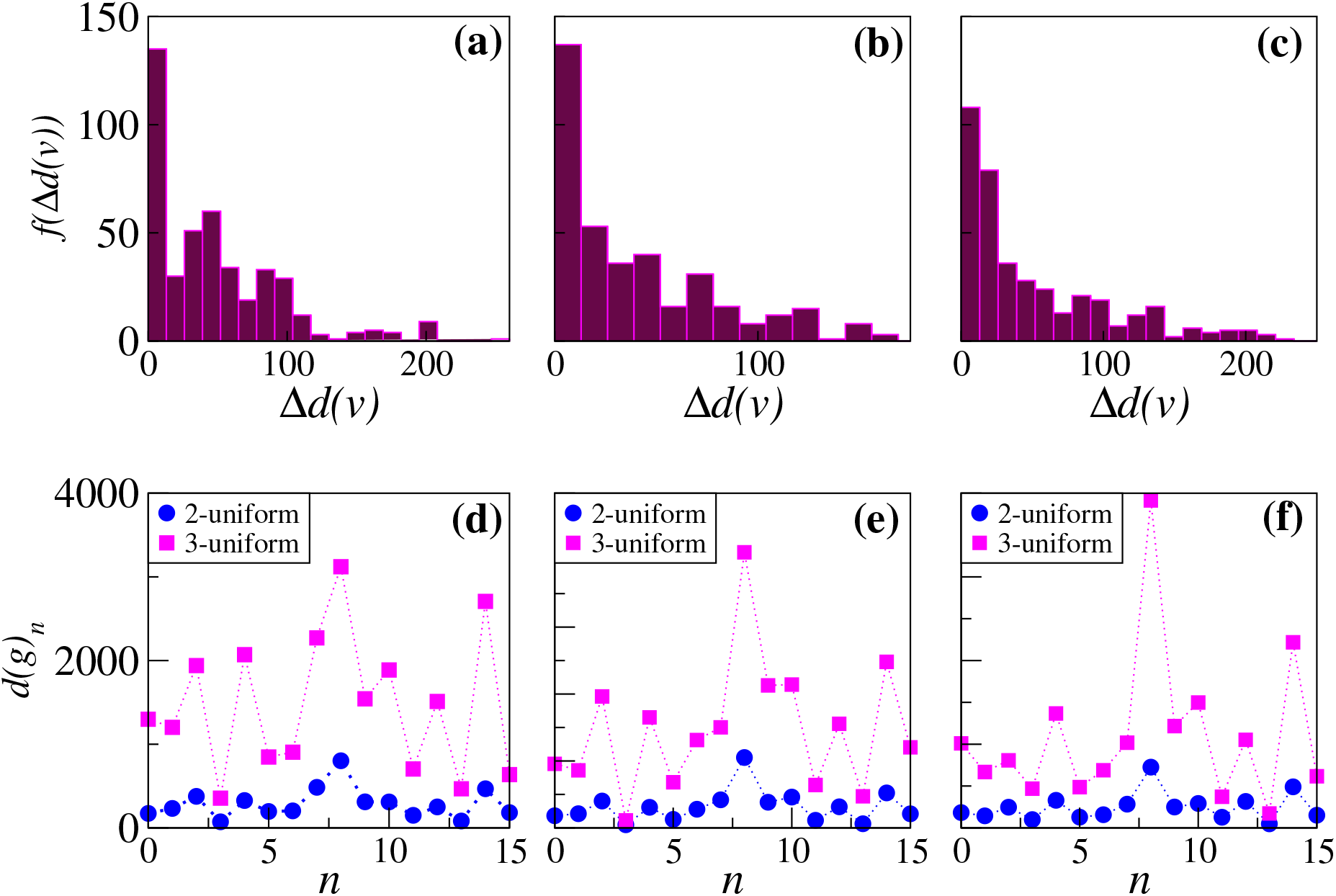
Distribution of the degree difference between 2- (*d*(*v*)_2_) and 3-uniform hyperedges (*d*(*v*)_3_) of variable sites, (Δ*d*(*v*)) for (a) low- (b) mid- and (c) high-altitude (upper panel). The 2- and 3-uniform hyperedge degrees (*d*(*g*)) for *n*^*th*^ gene plotted for (d) low-, (e) mid- and (f) high-altitude. The 3-uniform hyperedge degrees were found to be comparatively higher than the 2-uniform hyperedge degrees at all the altitudes (lower panel).

**Figure 4.**
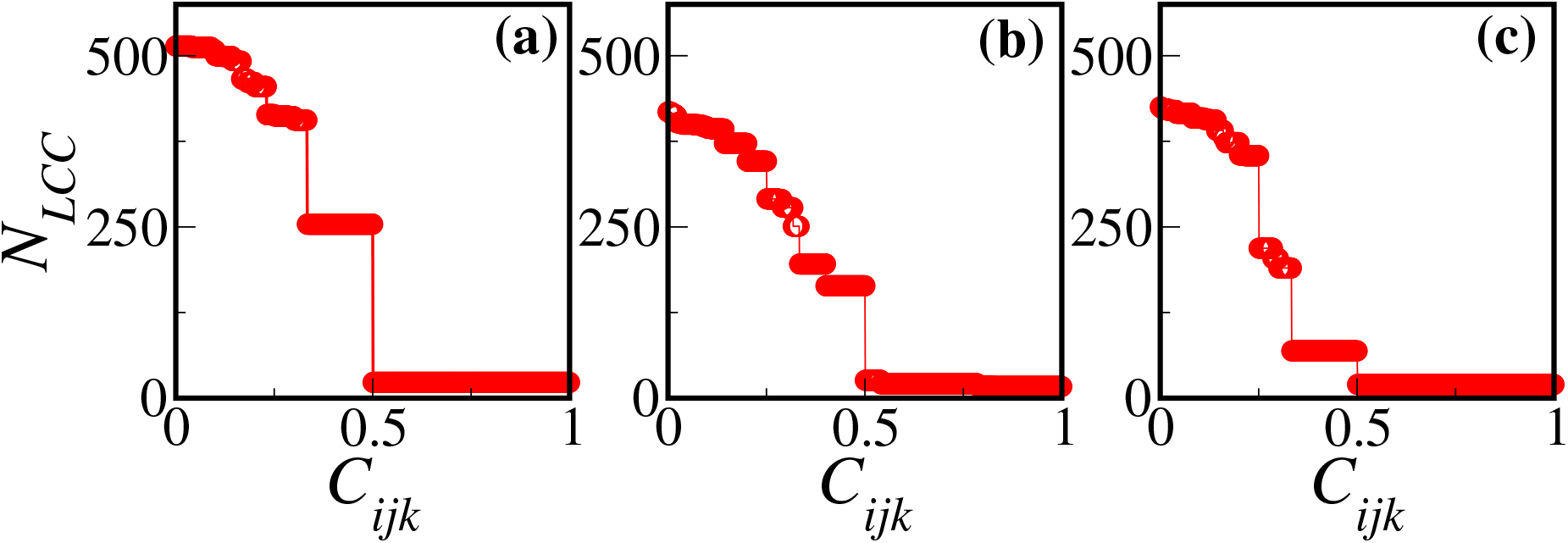
The size of the largest connected component (LCC) with respect to *C*_*ijk*_ to identify the threshold value for (a) low-altitude, (b) mid-altitude, and (c) high-altitude. There existed a sudden drop in the size of LCC at a particular *C*_*ijk*_, which we set as a co-mutation threshold (*C*_*th*_) to filter the hyperedges.

### Overlapping edges and associated genes

Next, we analyzed the statistics of the edge degree of each hypergraph. For 3-uniform hypergraphs, the edge degree was defined as the number of triangles a given edge is part of. After calculating the *d*(*v*)_*n*,3_, we mapped the edges (both nodes) to genes, yielding a gene pair for each edge. Since more than one edge can belong to the same gene pair, we get the gene-pair weight by adding the edge degree for all the edges of a given gene pair. For instance, edges (1,2) and (3,4) are part of two triangles each. Therefore, the edge degree of both these edges will be equal to 2. Next, mapping these edges to genes, both edges belong to the same gene-pair *G*1 − *G*2. Hence, the gene-pair weight will be equal to the sum of both the edge degrees, i.e., equal to four. This provided us with the information of overlapping edges between the triangles. In all the three altitude groups, *Control region* was commonly found to belong to hyperedges with the highest weights. Particularly, in low and mid-altitude, the control region formed the highest weight hyperedge with *ND5* gene and in high-altitude with *CYB* gene. Apart from the *Control region*, in low-altitude *ND5* formed highest weight hyperedge each with *ND1, ND2*, in mid-altitude *ND5* formed highest weight hyperedge each with *CO1, ND1, ND4*, and in high-altitude *CYB* formed highest weight hyperedge each with *ATP6, ND2, ND5*. In summary, in low-altitude ND gene yielded overlapping hyperedges. In high-altitude CYB, ATP6 and ND genes yield overlapping hyperedges. The CYB and ATP6 genes have already been established to play a significant role in high-altitude adaptation [40].

### Common and exclusive triangles

There existed few variable sites which were common and few others are exclusive among all the altitudes. We investigated how these variable sites form triangles by identifying the presence of shared and exclusive true triangles. Note that a variable site common to two altitudes could be part of two different triangles. Hence common variable sites are not necessarily part of common triangles for the corresponding altitudes. We found 181 common triangles among all the altitude groups. In addition, there are 123 common triangles between low- and mid-latitudes, 169 between low- and high-altitudes, and 612 between mid- and high-altitudes. A large proportion of triangles were found to be exclusive to each altitude group. We were interested in identifying the genes forming triangles in these common and exclusive triangles for each altitude. One simple method to access such genes is to map these common triangles to corresponding genes to get weighted gene triangles. The triangles with the highest weight had {*CYB* and *ND3*} genes present ubiquitously among common triangles of all the altitudes. Among common triangles of low- and mid-latitudes, {*ND2* and *ND4*} genes, and among common triangles of mid- and high-altitudes, {*CO1* and *ND5*} genes were present ubiquitously in gene triangles with the highest weights. To extend this in terms of genomics, the interaction between two particular genes was significant in forming triangles with a third different gene. This provided evidence and strengthened the presence of particular pair-wise interactions among mitochondrial genes.

### Codon positions in hyperedges

A variable site possesses information about codon position (CP) in the coding genes. The relation between co-evolving variable sites and codon positions has been described previously in the human population using two, and three-order motifs [39]. For a coding gene, the codon positions were set based on nucleotide position in codons 1, 2, and 3. For non-coding genes, all the nucleotide positions were set to 0. It was detected that CP 1 and 2 are highly conserved in the formation of the codon triangles, whereas CP 3 is versatile in the formation of these hyperedges (Fig. 5). The CP 3 predominates the formation of hyperedges with other coding and non-coding positions. The third codon position is considered to be weakly responsible for amino acid selection during protein synthesis, therefore, supporting the Wooble hypothesis it is not subjected to evolutionary constraint [41]. The codon-conservation has been observed in the regions of conopeptides revealing their accelerated evolution [42]. However, the hyperedges with all the associated sites being CP 3 are relatively less abundant than other hyperedges with one or two CPs occupied by coding and non-coding positions. This suggests that the third CP is subject to conformational stability provided by other CPs. Among all, CP 2 is the most conserved position, which supports the functional stability of amino acid selection during protein synthesis. Similarly, the hyperedges with all CP being 0 were also found to be relatively less. Since these do not code for any proteins, the associated hyperedges represent the intra-Control region and intra-RNA genes. This suggests that non-coding regions favour the formation of higher-order hyperedges least.

**Figure 5.**
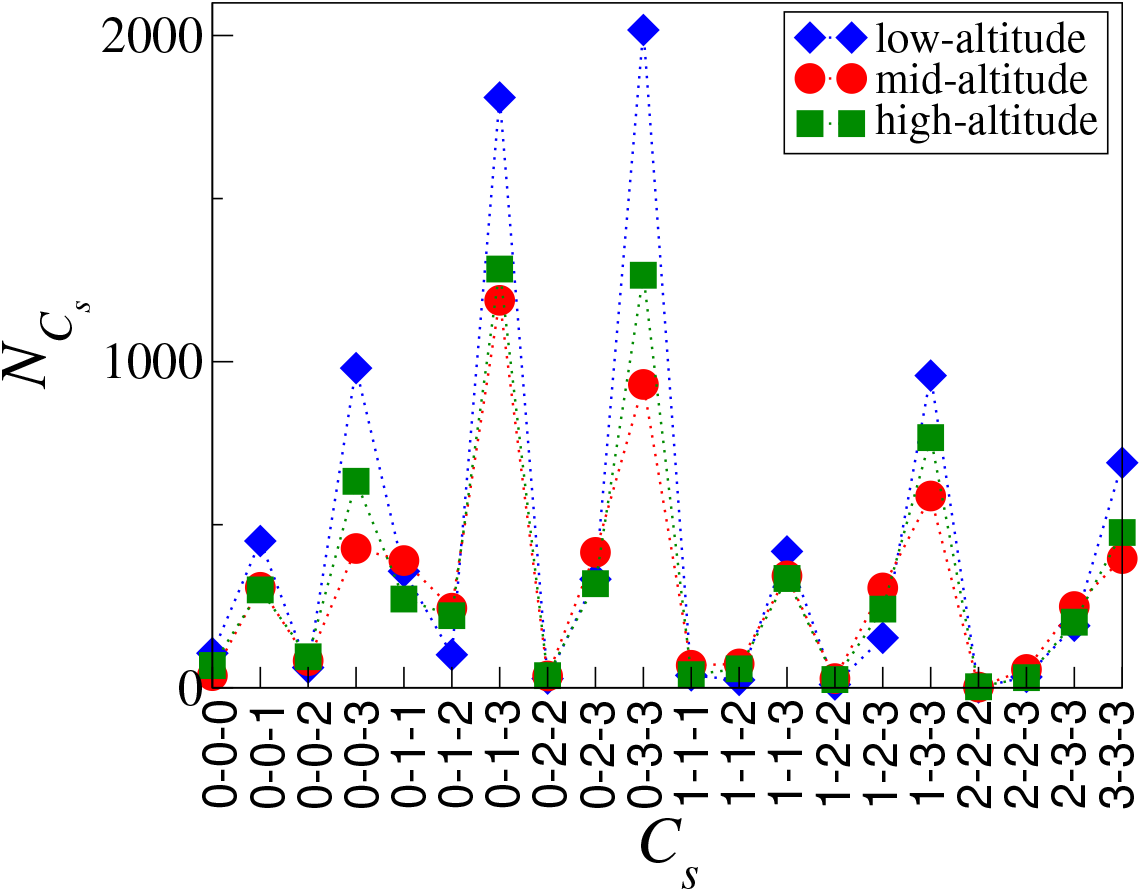
Distribution of codons in different altitude groups. *C*_*s*_ is codon type based on codon positions with *s* taking values 0, 1, 2 and 3, and 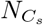 is *C*_*s*_ count.

### Gene triangles

After identifying the true triangles, we calculated the actual number of true gene triangles by summing over their occurrence in the network, thereby calculating their weights. If we exclude the *Control region* out of all the possible true gene triangles, only *∼*10% were identified as true gene triangles (Table 1) in all the altitudes. The *Control region* is known to be a highly mutating region of the mitochondrial genome. Owing to this, we excluded *Control region* to analyze the triangles of coding and non-coding genes explicitly. We plotted the distribution of weights (*p*(*W*(*g*))) of all the gene triangles and found that it follows the power-law (Fig. 6). This indicated that the mitochondrial genes prefer the formation of particular triangles over others due to the existence of conserved co-mutation patterns formed by variable sites. After selecting a specific *C*_*ijk*_ threshold value, all the variables sites of each surviving triangle were mapped to the corresponding genes, and the contribution of each gene was calculated as a relative weight for each altitude group (Fig. 7). The relative weight of each gene was defined based on the number and weight of the hyperedges for each gene. Note that for this calculation *Control region* variables sites were not considered. Note that *Control region* was a highly mutating region of mtDNA and hence occurred in all the groups equally. In the high-altitude group, *CYB* gene manifested distinctly high relative weight among coding genes and *tRNA-Gln* and *tRNA-Met* among non-coding genes (Fig. 7). In low and mid altitudes, none of the coding genes showed distinctly high relative weight; however, in the mid-altitude group, *tRNA-Ile, tRNA-Arg* and *tRNA-Leu*, and in the low-altitude group, *tRNA-Phe, tRNA-Ser* and *tRNA-Tyr* showed distinct relative weights among non-coding genes (Fig. 7). It was to note that in the low altitude, *tRNA-Phe* depicted distinctly high weight among non-coding genes; however, it was not found to be a high degree node.

**Figure 6.**
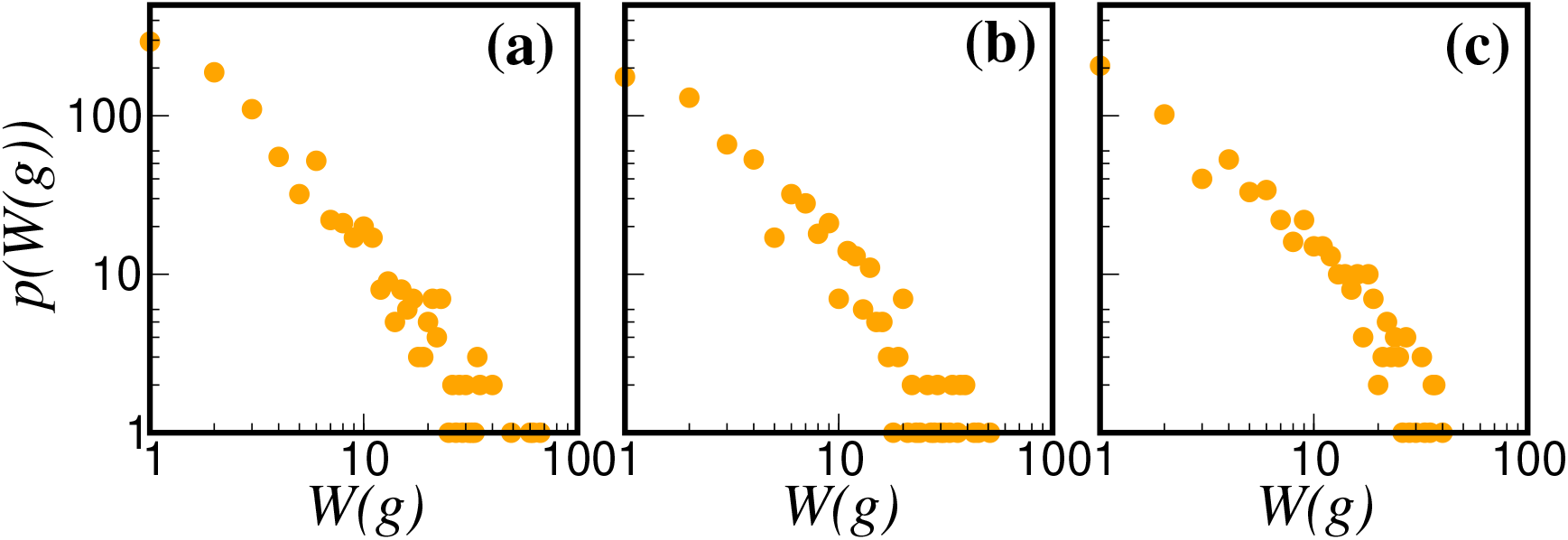
Weight distribution of gene triangles for (a) low-altitude, (b) mid-altitude, and (c) high-altitude.

**Figure 7.**
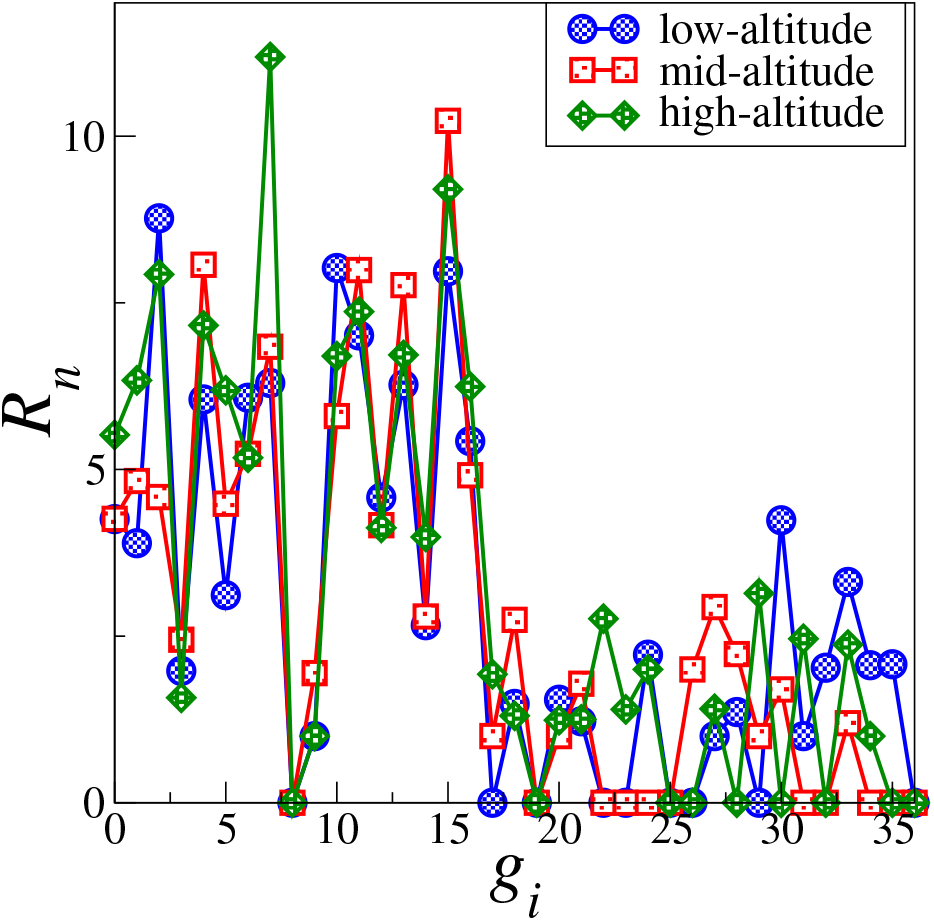
The relative weight (*R*_*s*_) of each gene for all three altitude groups. Note that Control region is excluded form this calculation.

### Gene hyperedges categories

Furthermore, we classified gene hyperedges into four categories based on the information of genes. These categories are *all-coding* (with no non-coding gene), *2-coding* (with two coding genes), *1-coding* (with one coding gene) and *0-coding* (with no coding genes). We found that *∼*50% of the gene triangles were from *2-coding* category, and *∼*1% from *all-coding* category in all the altitude groups. *All-coding and 1-coding* were present equally around 25%. The existence of a relatively high percentage of 2-coding type in all the altitudes suggested that the presence of one non-coding gene favoured the formation of higher-order interactions in the mitochondrial genome in general independent of environmental conditions. However, the gene triangles with high weights were different for different altitudes. The non-coding gene in the 2-coding category with high weight was found to be *12SrRNA* in low altitude and *16SrRNA* in both middle and high altitudes.

Primarily, we were interested in identifying specific gene triangles which might be important for high-altitude adaptation. To achieve this, we inspected for gene triangles having high weights (Table 2). From the table, it was clear that in low-altitude population *ATP6 and ND* genes, in mid-altitude population *CO1 and ND* genes, and in high altitude population *ATP6, CO1, CYB, and ND* genes contributed significantly in the formation of higher-order interactions. Earlier studies of perfectly co-occurring pair-wise interactions had revealed a similar set of genes with some additional genes for these different altitude groups. However, the variable sites forming gene triangles differed from those reported in Ref. [37]. This suggested that the formation of hyperedges was facilitated by a different set of variable sites, and might be capturing the genetic interactions according to their role in environmental adaptability. An important observation was that in each of the gene triangles, one of the three edges depicted considerable weight (discussed previously as edge weight), which was again observed when we inspected for the variable sites contributing to gene triangles with high weights. In the low-altitude group, ND1, ND5 genes or ATP6, ND5 genes corresponded to variable sites with the highest contribution, in mid-altitude, CO1, ND5 genes, and in high-altitude, CO1, ND1 or ATP6, CYB or ND5, CYB or ATP6, CO1 or ATP6, CYB genes in one or the other gene triangles corresponded to variable sites with the highest contribution. It was observed in the analysis of uniform hypergraphs for each altitude, specific genes were forming higher-weight triangles which assisted in further analysis at the phenotypic level.

## Conclusion

We investigated interaction hypergraphs of variable sites of mtDNA constructed based on the co-mutation frequency of three variable sites. Such hypergraphs were first filtered through the statistical score and second through gene-based information. We superimposed the variable site onto genes to finally realize the 3-uniform gene hypergraphs. Using the p-value test, *∼*10% higher-order interactions were found to be true hyperedges. These hyperedges belonged to distinct haplogroups in different altitude populations. At the low-altitude, they corresponded to M-haplogroup, in the mid-altitude, C-haplogroup, and in the high-altitude, the K-haplogroup. Moreover, codon-based hyperedges provided evidence about the codon bias and conservation of codon usage throughout the mitochondrial genome for all the altitude groups. Based on gene hyperedges, we found that in the low-altitude group, *ATP6 and ND* genes, in middle altitude group, *CO1 and ND5* genes, and in high altitude group, *CYB and ND5* genes were predominantly forming hyperedges. Here we have only considered 3-uniform hypergraphs, it would be interesting to investigate other higher-order interactions, say 4-, 5-uniform hyper to get further insights and to identify the hyperedges-based communities in the mitochondrial genome. Moreover, the analysis presented here provided a sophisticated method to pin down particular genes based on higher-order interactions and can be extended further to other biological systems composed of a large number of units and their interactions.

## Acknowledgments

SJ acknowledges SERB Power grant SPF/2021/000136, and RKV thanks UGC-CSIR for SRF (305089) fellowship. The work used the computational facility received from the Department of Science and Technology, Government of India, under FIST scheme (Grant No. SR/FST/PSI-225/2016) We thank Narayan Ganpati and Jerry David for their excellent comments throughout. SJ thanks Stefano Boccaletti for various useful suggestions during his visit to IIT Indore under VAJRA scheme (VJR/2019/000034). We are indebted to Mehraj Anvari, Mikhail Ivanchenko for useful comments on the final version of the manuscript.

